# Indoleamine 2,3-dioxygenase-1 expressing aggregate bone marrow dendritic cell populations are associated with systemic T-cell compartment changes in chronic myelomonocytic leukemia

**DOI:** 10.1101/2020.05.14.096297

**Authors:** Abhishek A. Mangaonkar, Kaaren K. Reichard, Moritz Binder, Giacomo Coltro, Terra L. Lasho, Ryan M. Carr, April Chiu, Vivian Negron, Mehrdad Hefazi, Theodora Anagnostou, Jose C Villasboas, Wilson Gonsalves, Naseema Gangat, Mithun Shah, Hassan B Alkhateeb, Aref Al-Kali, Michelle A Elliott, Kebede H Begna, Alexandra P Wolanskyj-Spinner, Mark R Litzow, William J Hogan, Stephen M Ansell, Animesh Pardanani, Ayalew Tefferi, Mrinal M. Patnaik

**Author notes:** **Corresponding author:** Mrinal M. Patnaik, MD, Associate Professor of Medicine, Division of Hematology, Department of Medicine, Mayo Clinic, Rochester, MN.

## Abstract

Systemic immune tolerance is not well-characterized in chronic myelomonocytic leukemia (CMML). Due to the presence of clonal plasmacytoid dendritic cells (pDC) in CMML, and the established association of lymph node indoleamine 2,3-dioxygenase-1 (IDO1)-positive (+) DC populations (IDC) with systemic immune tolerance in other malignant contexts, we sought to determine the association of IDO1 expression and bone marrow (BM) DC populations with systemic T-cell compartment changes using primary CMML patient samples (BM, plasma, and peripheral blood mononuclear cells) via immunohistochemistry (IHC), liquid chromatography-mass spectrometry (LC-MS), and time-of-flight mass cytometry (CyTOF). Our results highlight that aggregate BM IDC (CD123 and/or CD11c positive) occur in 33% CMML patients at any disease time-point (IHC), correlate with accentuated tryptophan catabolism (LC-MS, increased kynurenine level, median 4.7 versus 3 microM, P=0.049*), systemic regulatory T-cell expansion (CyTOF, %parent cell type, 14.5 versus 4.9%, P=0.04*) and play a role in disease progression, as evidenced by a higher rate of transformation to acute myeloid leukemia (41 versus 13%, P=0.002**), when compared to CMML patients without BM IDC. Our data also highlight a perturbed immune system in CMML with specific systemic immune signatures, particularly type 1, IL-17 producing helper T, CD4 terminal effector and natural killer cell suppression.

**Key Points:**

- Aggregate IDO1+ dendritic cell populations occur in the CMML bone marrow microenvironment, and their presence correlates with disease progression.
- Systemic immune microenvironment signatures in CMML indicate an altered T- and natural killer (NK)-cell balance. Specifically, suppression of type 1 helper T (Th1), IL-17 producing helper T (Th17), CD4 terminal effector and NK cells.
- IDO1+ bone marrow dendritic cell populations in CMML are associated with a T-cell compartment shift towards a regulatory T cell phenotype.

## Introduction

Chronic myelomonocytic leukemia (CMML) is a myelodysplastic/myeloproliferative neoplasm with an inflammatory mileu^1^ and a clinical association with autoimmune disorders/inflammatory syndromes ^2–4^, suggesting pervasive immune deregulation. Upregulation of specific immune checkpoints (PD-1, PD-L1 and PD-L2) in CD34+ isolated cells from patients with CMML compared to acute myeloid leukemia (AML) has been shown previously^5^; however, specific mechanisms of immune tolerance remain to be elucidated.

Numerous studies have reported the presence of plasmacytoid (p) dendritic cell (DC) populations in CMML bone marrow (BM)^6–8^, with a recent study concluding that these are clonal and associate with RAS pathway mutations.^8^ Previous studies using melanoma mouse models have established a role for indoleamine 2,3-dioxygenase-1 (IDO1) in regulating pDC^9–11^ and myeloid-derived (m) DC plasticity^12,13^, inducing tolerogenic phenotypes in both contexts. IDO1 is an immune checkpoint enzyme that induces systemic immune tolerance through multiple mechanisms, including regulatory T cell (Treg) expansion and tryptophan catabolism.^14,15^ In AML, IDO1 expression on blasts has identified as an independent adverse prognostic factor^16–18^, and shown to impair immune response by Treg induction.^19^ In CMML, the role of IDO1 in DC [both CD123+ pDCs and CD11c+ mDCs] and T-cell interactions, and impact on systemic immune microenvironment remains undefined.

## Methods

CMML patient samples [BM tissue blocks, peripheral blood mononuclear cell (PBMC), and plasma] from our clinically and molecularly-annotated biobank were obtained after regulatory approval.

Immunohistochemisty [IHC, Hematoxylin and Eosin (H&E), IDO1, CD123 and CD11c), multiplex immunostaining (IDO1, CD123 and CD11c), liquid chromatography mass spectrometry (LC-MS), time-of-flight mass cytometry (CyTOF), cytokine profiling (luminex array), and gene expression analysis (RNAseq) were performed in-house. IHC slides were independently reviewed by two hematopathologists. Detailed protocols and methods are available in the supplementary section.

### Aggregate IDO1+ DC populations in the BM microenvironment are associated with disease progression in CMML

IHC with H&E and IDO1 stains was performed on BM biopsies of 103 CMML patients (80 at diagnosis). At least one morphologically-defined BM IDO1+ DC population (IDC, consensus agreement by two hematopathologists, 90% concordance) was identified in 34 (33%) patients (25% at diagnosis). DC phenotypic characterization was done by CD123 and CD11c staining (Figure 1A-1D), with 18 (56%) being positive for both, 7 (22%) for CD123, and 5 (16%) for CD11c only (2-negative for both, 2-status unknown) suggesting that IDO1 expression is not limited to the previously described CD123-marked pDC populations in CMML^8^. IDO1 expression in the DCp vicinity was further confirmed by multiplex (IDO1, CD123 and CD11c) staining (Figure 1E). In patients without IDC at diagnosis (n=60), 1 (2%) acquired them at disease progression after hypomethylating agent therapy, while 2 (3%) at AML transformation. This indicates that BM IDC play a role in CMML disease progression, a conclusion further strengthened by the observed higher frequency of AML transformation (41% versus 13%, P=0.002**) and an adverse Kaplan-Meier estimate of AML-free survival (median, 122 versus 107 months, P=0.04*, Figure 1F, supplementary table 1) in patients who acquired these aggregates at any time-point, despite minimal differences in clinical, genetic and prognostic characteristics (supplementary table 1). Unlike in AML where IDO1 expression is seen in leukemic blasts^16–18^; in CMML, IDO1 is expressed in aggregate DC populations in the BM microenvironment and correlates with disease progression.

**Figure 1:**
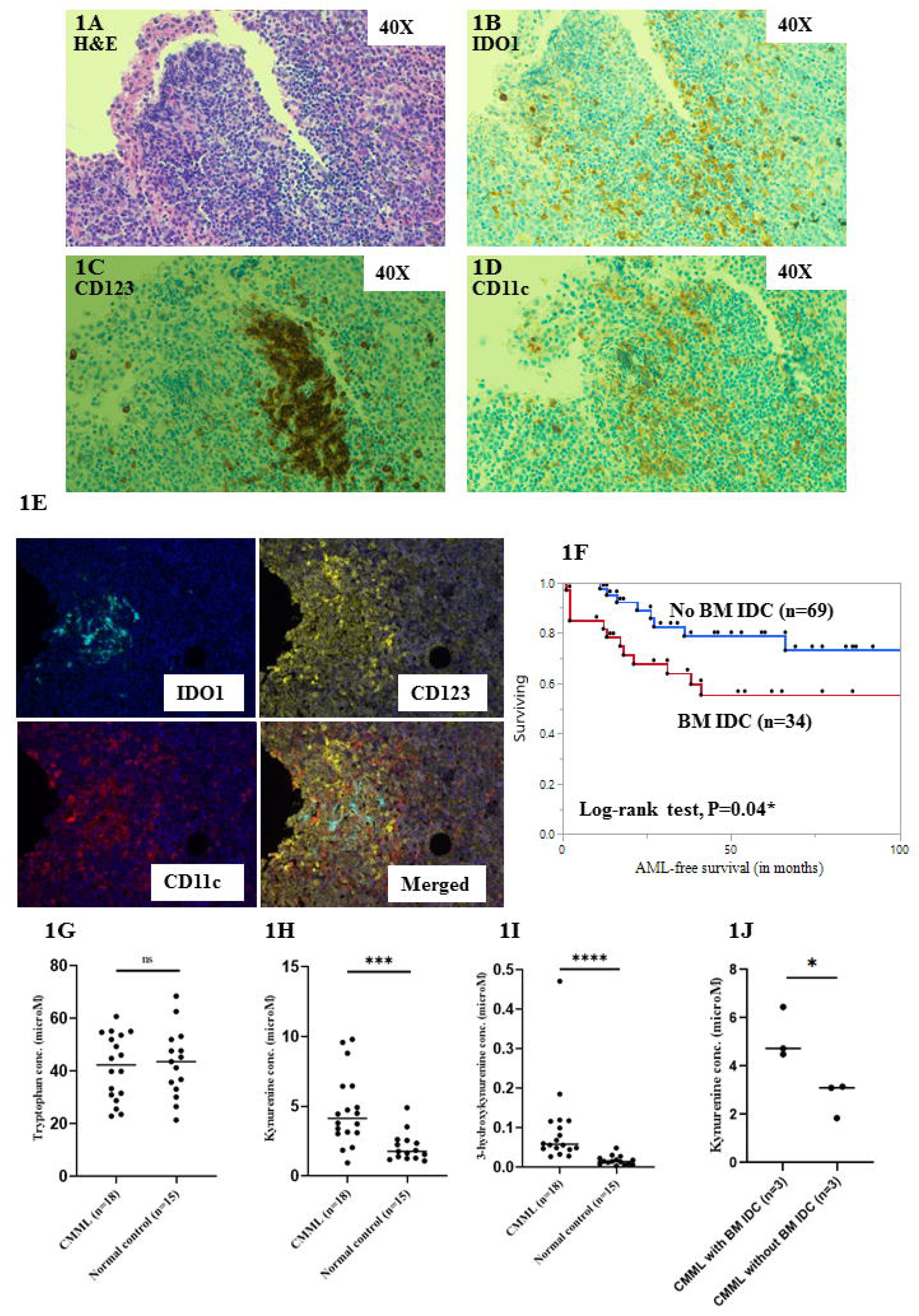
Figure shows that indoleamine 2,3-dioxygenase-1 (IDO1)-positive dendritic cell populations (IDC) are present in the chronic myelomonocytic leukemia (CMML) bone marrow (BM) microenvironment, associate with adverse acute myeloid leukemia (AML)-free survival, and associate with metabolic signatures of systemic IDO1 activity. (1A) shows hematoxylin and eosin (H&E) staining of an area of CMML BM with IDC (40X magnification), (1B) shows IDO1 staining (40X magnification), (1C) shows CD123 staining, and (1D) shows CD11c staining. (1E) shows multiplex immunostaining showing clustering of IDO1 expression with CD123 and CD11c stains in a CMML BM, confirming expression in a DC population. (1F) shows Kaplan-Meier estimate of AML-free survival in CMML patients with IDC at any disease time-point (n= 33) versus CMML patients without IDC (n=69, median 122 versus 107 months, log-rank P value = 0.04*). (1G), (1H) and (1I) shows similar tryptophan (42.4 versus 43.6 microM, P=0.9), higher kynurenine (4.1 versus 1.8 microM, P=0.0006***), higher 3-hydroxykynurenine (0.06 versus 0.01 microM, P<0.0001****) concentrations in CMML patients (n=18) and normal controls (n=15) respectively, as evaluated by liquid chromatography-mass spectrometry (LC-MS) in plasma samples. (1J) shows higher kynurenine concentration in diagnostic plasma samples (n=6) of CMML patients with versus without BM IDC (4.7 versus 3 microM, P=0.04*).

### Metabolic signatures of systemic IDO1 activity are evident in CMML

We then assessed for evidence of systemic IDO1 activity in CMML by measuring tryptophan and its metabolites (kynurenine and 3-hydroxykynurenine). Through LC-MS, although median tryptophan concentrations (42.4 versus 43.6 microM, P=0.9, figure 1G) were not significantly different between CMML (n=18) and normal control plasma samples (n=15), kynurenine (4.1 versus 1.8 microM, P=0.0006***, figure 1H), and 3-hydroxykynurenine (0.06 versus 0.01 microM, P<0.0001****, figure 1I) concentrations were higher in CMML (supplementary table 2). These findings suggest accentuated tryptophan catabolism in CMML, a signature of systemic IDO1 activity.^20^ Further, median kynurenine concentration was higher in CMML patients with versus without BM IDC at diagnosis (4.7 versus 3 microM, P=0.049*, figure 1J), suggesting an increased systemic IDO1 activity in the former group.

### CyTOF assessment highlights an altered T and natural killer (NK) cell balance in CMML

In order to characterize the immune microenvironment in CMML, we selected untreated (except hydroxyurea) CMML PBMC (n=7, supplementary table 1) samples and age-matched normal PBMC controls (n=3), and performed CyTOF analysis. Quantitative comparisons [percentage (%) of parent cell type], indicated significant differences between CMML and controls including a decreased type 1 helper T-cells (2.4 versus 10.8, p=0.03*), Th1/type 2 helper T-cell (Th2) ratio (0.2 versus 0.7, p=0.02*), IL-17 producing T-helper (Th17) cells (0.5 versus 7.3, p=0.02*), CD4 terminal effector cells (11.4 versus 23.2, P=0.03*), and NK cells [2.2 versus 17.4, P=0.02*, (supplementary table 2 and figure 2A-2E)]. Suppression of NK, Th17, and Th1 cells has been shown in AML^21,22^, and CML^23^, suggesting similar patterns of immune dysregulation in CMML.

**Figure 2:**
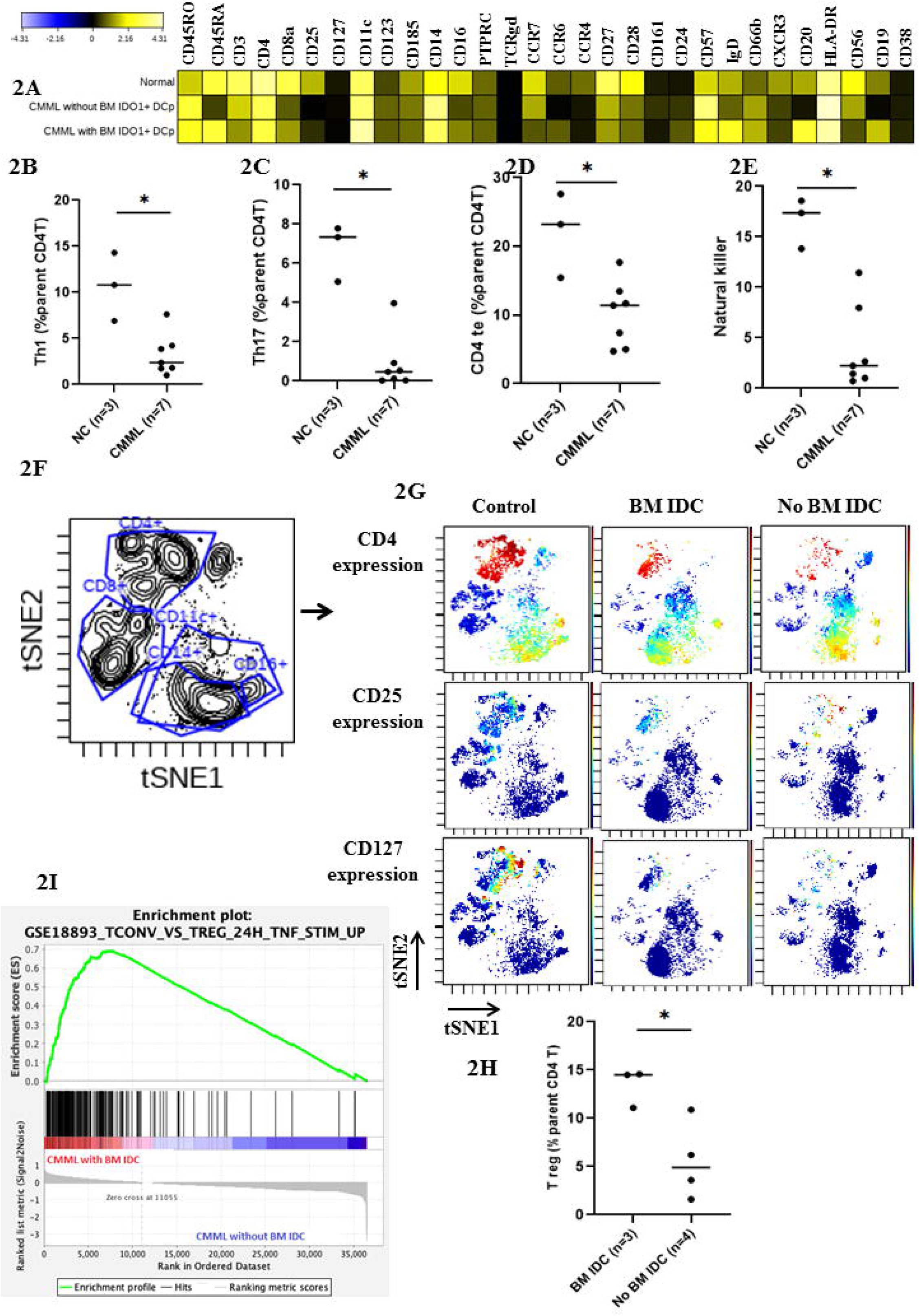
Figure 2 shows mass cytometry (CyTOF) analysis of peripheral blood mononuclear cell (PBMC) samples from untreated patients with chronic myelomonocytic leukemia (CMML) and age-matched normal controls. (2A) shows a heatmap of all the markers used in the CyTOF panel (details in supplementary) and stratified by a representative sample in each group [normal control, CMML without bone marrow (BM) indoleamine 2,3-dioxygenase-1 (IDO1) dendritic cell populations (IDC), and CMML with IDC]. Expression is indicated as transformed ratio of means. (2B), 2(C), 2(D), and 2(E) shows a significantly suppressed median percentage (%) of parent cell populations of type 1 helper T cells (Th1, median 2.4 versus 10.8, P=0.03*), IL-17 producing helper T cells (Th17, 0.5 versus 7.3, P=0.02*), CD4 terminal effector (te) cells (11.4 versus 23.2, P=0.03*) and natural killer (NK, 2.2 versus 17.4, P=0.02*) cells in CMML versus normal controls respectively. (2F) shows visualization of dimensionality reduction analysis (viSNE) plot with gates delineating CD4+, CD8+, CD11c+, Cd14+, and CD16+ populations. (2G) shows expression of specific markers such as CD4, CD25 and CD127 (red denotes increased expression). This figure provides a visualization of an expanded regulatory T cell (Treg, CD4+/CD25+/CD127^dim^) population in a CMML patient with versus without BM IDC. (2H) shows quantitative analysis confirming an expanded T reg (%parentCD4 T cell) population in untreated CMML patients with BM IDC (n=3) versus untreated CMML patients without BM IDC (n=4, median 14.5 versus 4.9, P=0.04*). (2I) shows gene set enrichment analysis showing significant enrichment of T reg associated genes (normalized enrichment score = 3.97, p<0.0001***) through RNA-seq analysis of untreated PBMC samples derived from CMML patients (n=4, indicated in red) with versus without (n=4, indicated in blue) BM IDC (samples collected at the same time-point as IHC assessment).

### Aggregate IDO1^+^ bone marrow DC populations are associated with expanded systemic regulatory T cell compartment in CMML

When untreated CMML patients with (n=3) and without BM IDC (n=4) were compared, the former group showed higher percentage (%, of parent cell type) T regs (14.5 versus 4.9%, P=0.04*, supplementary table 3, figure 2F-2H). This suggests that IDC in CMML BM microenvironment promotes Treg differentiation in the systemic T-cell compartment. As shown in the context of other maligancies^9^, this is likely due to DC-T-cell interactions driven by T-cell recruitment to these locations. This was confirmed by cytokine profiling which showed higher median levels (pg/ml) of RANTES (CCL5, 8650 versus 349.7, p=0.04*) and MIP-1β (macrophage inflammatory protein-1β, 28.7 versus 10.8, P=0.049*), known T-cell chemoattractants^24,25^, in diagnostic plasma samples of CMML patients with BM IDO1+ DCp (supplementary table 4). Further, in diagnostic samples (plasma and PBMC, n=5) collected at the same time-point, kynurenine levels showed a significant strong positive correlation with % T-reg populations (Spearman rho=0.9, P=0.04*, supplementary figure 1). Additional confirmation was performed with gene set enrichment analysis in a subset of PBMC RNA derived from untreated patients at the time of IHC assessment which indicated T reg associated gene upregulation (normalized enrichment score = 3.97, p<0.0001***) in untreated CMML patients with versus without BM IDC (n=4 each, figure 2H).

In summary, our constellation of immunohistochemical, LC-MS, CyTOF, cytokine profiling and gene set enrichment results provide evidence of systemic immune dysregulation in CMML, and highlight the association of bone marrow IDO1+ DC populations with a T cell compartment shift towards a regulatory T cell phenotype. Future research is needed to clarify the genetic/epigenetic events responsible for CMML IDO1+ DCp formation, to pinpoint the biology of DC-T cell interactions, recapitulate the findings in *in-vivo* models and explore therapeutic vulnerabilities with IDO1 and/or CD-123 directed therapies.

## Acknowledgements

Authors would like to acknowledge the services and expertise of Pathology Research Core, Immune Monitoring Core, and Metabolomics Resource Core of Mayo Clinic, Rochester, MN. This work was supported in parts by the Conquer Cancer Foundation of American Society of Clinical Oncology-sponsored Young Investigator Award and Mayo Clinic Hematology Small Grants Program Grant awarded to AAM and Grant #UL1TR002377 from the National Center for Advancing Translational Sciences (NCATS) awarded to MMP. Results and views do not necessarily reflect the views of the funding agencies.

## Author contributions

AAM co-designed the study with MMP, performed key experiments, data analysis, and wrote the first draft of the manuscript. GC, TLL, RC, MB, performed key experiments and provided critical feedback. AC, VN and KKR reviewed hematopathology slides and provided critical feedback. TA, JCV, MH and SMM provided feedback on mass cytometry experiment and data interpretation. WG aided mass spectrometry experiment and data analysis. NG, AP, AT, WG, MRL, KHB, HA, MS, APW, and WH contributed patient samples and edited the manuscript. MMP supervised the study, and edited all manuscript drafts. All authors participated in writing and editing the manuscript.

