## Supplementary file for "Indoleamine 2,3-dioxygenase-1 expressing aggregate bone marrow dendritic cell populations are associated with systemic T-cell compartment changes in chronic myelomonocytic leukemia"

***Supplementary Table 1:* Table displaying distribution of variables for all chronic myelomonocytic leukemia (CMML) patients with and without indoleamine 2,3-dioxygenase-1 positive dendritic cell populations (IDC) in the bone marrow (BM) microenvironment at any time-point in disease diagnosis.**

| ***Variable; Median value (range or %)*** | ***All (n=103)*** | ***BM IDC (n=34)*** | ***No BM IDC***  ***(n=69)*** | ***P value*** |
| --- | --- | --- | --- | --- |
| Age (years) | 70 (33-91) | 69 (44-91) | 70 (33-87) | 0.6 |
| No. of males; | 67 (66) | 21 (62) | 46 (69) | 0.5 |
| Hb; gm/dl | 10.9 (6.7-15.4) | 11.3 (6.7-15.4) | 10.8 (6.8-15.1) | 0.6 |
| WBC count x 10^9^ per liter | 12.4 (2-185.7) | 12.9 (2-185.7) | 12.4 (3.8-126.2) | 0.8 |
| Platelet count x 10^9^ per liter | 101 (11-726) | 102.5 (24-726) | 101 (11-383) | 0.9 |
| AMC x 10^9^ per liter | 2.4 (1-40) | 2.5 (1-29.7) | 2.4 (1-40) | 0.5 |
| BM blasts | 2 (0-19) | 2 (0-19) | 2 (0-15) | 0.1 |
| PB blasts | 0 (0-19) | 0 (0-19) | 0 (0-18) | 0.08 |
| Autoimmune features* | 38 (37) | 16 (47) | 22 (32) | 0.1 |
| *2016 WHO classification (Evaluable=101)* | | | | |
| CMML-0 | 67 (66) | 18 (53) | 49 (73) | 0.08 |
| CMML-1 | 17 (17) | 8 (24) | 9 (13) |  |
| CMML-2 | 17 (17) | 8 (24) | 9 (13) |  |
| *FAB classification (Evaluable=100)* | | | | |
| Dysplastic | 55 (55) | 19 (58) | 36 (54) | 0.7 |
| Proliferative | 45 (45) | 14 (42) | 31 (46) |  |
| Abnormal cytogenetics (Evaluable=101) | 34 (34) | 11 (32) | 23 (34) | 0.6 |
| *Next generation (n=91, panel details below**)* | | | | |
| Histone modification | | | | |
| *ASXL1* | 53 (58) | 19 (58) | 34 (59) | 0.9 |
| *EZH2* | 3 (3) | 3 (9) | - | **0.01** |
| DNA methylation | | | | |
| *TET2* | 32 (35) | 13 (39) | 19 (33) | 0.5 |
| *DNTM3A* | 6 (7) | 1 (3) | 5 (9) | 0.3 |
| Dual epigenetic effect | | | | |
| *IDH1* | - | - | - | - |
| *IDH2* | 6 (7) | 4 (12) | 2 (3) | 0.1 |
| Splicing | | | | |
| *SRSF2* | 47 (53) | 18 (58) | 29 (50) | 0.5 |
| *SF3B1* | 2 (2) | - | 2 (3) | 0.2 |
| *ZRSR2* | 3 (3) | 1 (3) | 2 (3) | 0.9 |
| *U2AF1* | 9 (10) | 4 (12) | 5 (9) | 0.6 |
| Cytokine signaling | | | | |
| *JAK2* | 4 (4) | 2 (6) | 2 (3) | 0.6 |
| *NRAS* | 14 (16) | 6 (19) | 8 (14) |  |
| *CBL* | 20 (22) | 9 (27) | 11 (19) |  |
| *MPL* | 1 (1) | - | 1 (2) | 0.3 |
| *CALR* | - | - | - | - |
| *KRAS* | 7 (8) | 2 (6) | 5 (9) | 0.7 |
| *KIT* | 2 (2) | - | 2 (3) | 0.2 |
| *PTPN11* | 5 (5) | 2 (6) | 3 (5) | 0.9 |
| *SH2B3* | 1 (1) | - | 1 (2) | 0.3 |
| Signal transduction/transcription/metabolism | | | | |
| *RUNX1* | 6 (7) | 4 (12) | 2 (4) | 0.1 |
| *SETBP1* | 16 (18) | 5 (15) | 11 (19) | 0.6 |
| *FLT3* | 3 (3) | 3 (9) | - | **0.01** |
| *BCOR* | - | - | - | - |
| *CEBPA* | 1 (1) | - | 1 (2) | 0.3 |
| *CSF3R* |  | - | - | - |
| *NPM1* | - | - | - | - |
| Tumor suppressor | | | | |
| *TP53* | 3 (3) | 2 (6) | 1 (2) | 0.3 |
| *Disease prognostication per contemporary models* | | | | |
| Mayo Prognostic Model (Evaluable=100) | | | | |
| Low | 38 (38) | 10 (30) | 28 (42) | 0.5 |
| Intermediate | 31 (31) | 11 (33) | 20 (30) |  |
| High | 31 (31) | 12 (36) | 19 (28) |  |
| Mayo Molecular Model (Evaluable=90) | | | | |
| Low | 6 (7) | 3 (9) | 4 (6) | 0.5 |
| Intermediate-1 | 27 (30) | 7 (22) | 24 (37) |  |
| Intermediate-2 | 29 (32) | 11 (34) | 19 (29) |  |
| High | 28 (31) | 11 (34) | 18 (28) |  |
| Groupe Français des Myélodysplasies (GFM, evaluable=97) | | | | |
| Low | 39 (40) | 14 (44) | 25 (38) | 0.9 |
| Intermediate | 45 (46) | 14 (44) | 31 (48) |  |
| High | 13 (13) | 4 (13) | 9 (14) |  |
| Spanish cytogenetic risk stratification (Evaluable=100) | | | | |
| Low | 71 (71) | 23 (70) | 48 (72) | 0.9 |
| Intermediate | 16 (16) | 5 (15) | 11 (16) |  |
| High | 13 (13) | 5 (15) | 8 (12) |  |
| ***Outcomes*** | | | | |
| Median follow-up (95% CI) | 87 (53, 196) | 54 (27, NR) | 196 (66, 196) | 0.5 |
| Transformation to AML | 23 (23) | 14 (41) | 9 (13) | **0.002** |
| AML-free survival (95% CI) | 122 (107, NR) | 122 (21, NR) | 107 (107, NR) | **0.04** |
| Overall survival (95% CI) | 34 (21, 46) | 38 (21, 65) | 29 (16, 46) | 0.5 |

**Abbreviations:** No.=Number; Hb=hemoglobin, WBC=white blood cell; AMC=absolute monocyte count; BM=bone marrow, PB=peripheral blood, NR=not reached, CI=Confidence Interval, AML=acute myeloid leukemia

Statistically significant P values are indicated in bold.

*Definition of autoimmune features was used as per standard clinical diagnostic criteria as mentioned in ref. 4.

**Two panels were used at diagnosis or first referral with details as below (the research NGS panel was used in cases where clinical NGS was not performed).

| **Institution** | **Gene panel** | **List of genes** | **Coding region coverage** | **Read depth** |
| --- | --- | --- | --- | --- |
| **Mayo**  **Clinic** | OncoHeme NGS  for Hematologic Cancers | *ASXL1; BCOR; BRAF; CALR; CBL; CEBPA; CSF3R; DNMT3A; ETV6; EZH2; FLT3; GATA1; GATA2; IDH1; IDH2; JAK2; KIT; KRAS; MPL; MYD88; NOTCH1; NPM1; NRAS; PHF6; PTPN11; RUNX1; SETBP1; SF3B1; SRSF2; TERT; TET2; TP53; U2AF1; WT1; ZRSR2* | Variable per gene.  See details at <https://www.mayomedicallaboratories.com/test-catalog/Overview/63367> | >250X |
|  | Research NGS | *ASXL1; ASXL2; ATM; BCOR; BCORL1; CALR; CBL; CEBPA; CSF3R; DNMT3A; EED; ETNK1; EZH2; FLT3; GATA2; IDH1; IDH2; JAK2; JARID2; KIT; KRAS; MPL; NRAS; PHF6; PTPN11; RPS6KA2; RUNX1; SETBP1; SF3B1; SH2B3; SRSF2; STAG2; STK11; SUZ12; TERC; TERT; TET1; TET2; TET3; TP53; U2AF1; ZRSR2* | Full exon region coverage for each gene. | >500X |

***Supplementary Table 2:* Table displaying distribution of tryptophan and downstream metabolites (kynurenine, 3-hydroxykynurenine) in normal controls, and chronic myelomonocytic leukemia (CMML) patients with and without indoleamine 2,3-dioxygenase-1 positive dendritic cell populations (IDC) in the bone marrow microenvironment at disease diagnosis as assessed by liquid chromatography-mass spectrometry (LC-MS, protocol details below).**

| ***Variable; Median value (range)*** | **Controls (n=15)** | **CMML (n=18)** | **LC-MS done at CMML diagnosis**  **(n=6)** | **LC-MS not done at CMML diagnosis**  **(n=12)** | ***P value (Controls versus CMML)** | ****P value (CMML with versus without IDC at diagnosis)** |
| --- | --- | --- | --- | --- | --- | --- |
| Tryptophan (µM) | 43.6  (21.4-68.5) | 42.4  (22.9-61) | 40.5  (22.9-60.7) | 42.4  (23.6-55.1) | 0.9 | 0.5 |
| Kynurenine (µM) | 1.8 (1.1-5) | 4.1  (0.9-9.8) | 3.8 (1.8-6.5) | 4.1 (1-9.8) | **0.0006** | **0.049** |
| 3-hydroxykynurenine (µM) | 0.01  (0.004-0.05) | 0.06  (0.03-0.5) | 0.05  (0.03-0.2) | 0.07  (0.03-0.5) | **<0.0001** | 0.5 |
| Tryptophan/kynurenine ratio | 23.4  (5.4-34.3) | 10.4  (3.4-58) | 12.5  (3.5-15.7) | 9.2  (3.4-58) | **0.0001** | 0.3 |
| Kynurenine/3-hydroxykynurenine ratio | 124.2  (45.2-777.3) | 63.8  (9.4-119.9) | 74.2  (32.7-95.2) | 62.1  (9.4-119.9) | **0.002** | 0.5 |

**Statistically significant P values are indicated in bold. **For calculation of this P value, only cases where the LC-MS was done at diagnosis (n=6) and had immunohistochemistry data were considered (three patients had BM IDC, while three did not).*

***Supplementary Table 3*:** **Table showing differences in mass cytometry (CyTOF)-determined median percentages of parent (Italicized cell types are the parent cell populations for the respective cell types below) cell populations (range) between patients with chronic myelomonocytic leukemia (CMML) and age-matched normal controls.**

| **Cell type** | **Normal (n=3)** | **CMML (n=7)** | **CMML without IDC (n=4)** | **CMML with IDC (n=3)** | **P value (Normal versus CMML)** | **P value (CMML without or with IDC)** |
| --- | --- | --- | --- | --- | --- | --- |
| CD3 T cells | 64.3 (56.7-68.6) | 69.8 (21.2-96.5) | 76.2 (68.2-83.3) | 56.7 (21.2-96.5) | 0.3 | 0.5 |
| *CD8 T cells* | 0.3 (0.2-31.4) | 0.42 (0-44.31) | 0.5 (0-44.3) | 0.2 (0.06-0.8) | 0.9 | 0.7 |
| *CD8 T cells subpopulations* | | | | | | |
| Naïve T cells | 13.9 (4.6-17.3) | 0.62 (0-11.8) | 1.1 (0-6.6) | 0 (0-11.8) | 0.05 | 0.7 |
| CD8 central memory cells | 0.9 (0.4-4.2) | 0.2 (0-20) | 1.5 (0-3.4) | 0.2 (0.02-20) | 0.4 | 0.5 |
| CD8 effector memory cells | 34.8 (7.9-48.9) | 20 (0-78.8) | 22.1 (0-78.8) | 20 (3.5-64.7) | 0.7 | 1.0 |
| CD8 terminal effector cells | 50.6 (30.2-86.5) | 60 (0-96.37) | 41.1 (0-79.3) | 60 (23.5-96.4) | 0.7 | 0.5 |
| *CD4 T cells* | 63.5 (56.4-70.2) | 57 (47.2-82.2) | 69.3 (47.2-82.2) | 52.1 (51-57) | 0.7 | 0.3 |
| *CD4 T cell subpopulations* | | | | | | |
| Regulatory T cells | 17 (8.8-19.4) | 10.9 (1.6-14.6) | 4.9 (1.6-10.9) | 14.5 (11.1-14.6) | 0.1 | **0.04** |
| T helper cells-1 | 10.8 (6.9-14.3) | 2.4 (1-7.6) | 1.8 (1-4.2) | 3.9 (2.4-7.6) | **0.03** | 0.2 |
| T helper cells-2 | 16.3 (14.9-17.5) | 17.1 (5.1-28.5) | 8.3 (5.1-28.5) | 20.1 (17.1-22.6) | 0.9 | 0.3 |
| Th1/Th2 ratio | 0.7 (0.4-0.9) | 0.2 (0.1-0.4) | 0.2 (0.1-0.4) | 0.2 (0.1-0.3) | **0.02** | 1.0 |
| Th17 | 7.3 (5.1-7.8) | 0.5 (0.03-4) | 0.1 (0.03-0.5) | 0.9 (0.5-4) | **0.02** | 0.08 |
| Naïve CD4 T | 17.7 (13.4-25.1) | 35.8 (15.5-51) | 43 (15.5-51) | 25 (23.2-35.8) | 0.1 | 0.3 |
| CD4 Central memory cells | 29.8 (26.9-37.9) | 28.1 (21.1-52.2) | 31.2 (25.3-52.2) | 24.2 (21.1-47.3) | 0.7 | 0.3 |
| CD4 Effector memory cells | 29.4 (21.1-32.6) | 19.9 (6.9-42.7) | 13.6 (7-42.7) | 22.2 (17.7-42.6) | 0.4 | 0.5 |
| CD4 Terminal effector cells | 23.2 (15.4-27.6) | 11.4 (4.7-17.7) | 6.2 (4.7-13.5) | 11.7 (11.4-17.7) | **0.03** | 0.2 |
| Gamma Delta T cells | 1.2 (0.3-3.6) | 0.4 (0.1-1.5) | 0.5 (0.2-1.5) | 0.3 (0.1-0.7) | 0.3 | 0.5 |
| B cells | 10.3 (9.2-14.5) | 8.1 (0.8-68.2) | 8.2 (1.3-21.2) | 2.5 (0.8-68.2) | 0.3 | 0.7 |
| *B cell subpopulations* | | | | | | |
| Naïve B cells | 74.3 (71.7-74.8) | 69.4 (39.7-95.3) | 79.5 (46.2-90.6) | 68.7 (39.7-95.3) | 0.7 | 0.7 |
| Memory B cells | 25.8 (25.2-28.3) | 30.6 (4.7-60.3) | 20.5 (9.4-54) | 31.3 (4.7-60.3) | 0.7 | 0.7 |
| Transitional B cells | 7.2 (5.8-23.4) | 2.2 (1.2-16.5) | 6.1 (1.2-16.5) | 2.2 (1.5-5.1) | 0.1 | 1.0 |
| Plasmablasts | 0.06 (0.03-0.2) | 0.2 (0-3.1) | 0.2 (0-0.64) | 0.4 (0-3.1) | 0.3 | 0.6 |
| NK cells | 17.4 (13.8-18.6) | 2.2 (0.7-11.5) | 2 (0.7-8) | 2.2 (1-11.5) | **0.02** | 0.7 |
| Monocytes | 29.3 (27.8-35.3) | 60 (18.4-87.8) | 54 (18.4-88) | 60 (53.2-69.8) | 0.08 | 1.0 |
| Dendritic cells | 1.3 (1.1-6.6) | 4.2 (0.6-22) | 8.9 (1.6-22) | 4.2 (0.6-6.5) | 0.4 | 0.6 |
| pDC | 49 (4.6-52.2) | 48.9 (18.5-96.4) | 34.6 (18.5-96.4) | 72.2 (29-89) | 0.6 | 0.6 |
| mDC | 51 (47.8-95.4) | 51.1 (3.6-81.5) | 65.4 (3.6-81.5) | 27.8 (11.1-71) | 0.6 | 0.5 |

***Abbreviations:*** h=helper; Reg=regulatory; NK=natural killer; pDC=plasmacytoid dendritic cells; mDC=myeloid dendritic cells.

| **Cytokine;**  **median values in pg/ml (range)** | **CMML**  **(n=6)** | **CMML**  **with**  **BM IDC (n=3)** | **CMML**  **without**  **BM IDC (n=3)** | **P value** |
| --- | --- | --- | --- | --- |
| EGF | 14  (0-71.3) | 15.4 (7.9-71.3) | 12.7 (0-17.5) | 0.5 |
| Eotaxin | 32.7  (20-67.3) | 26.1 (20-27.8) | 67.3 (37.7-67.3) | 0.05 |
| FGF-basic | 0  (0-2.6) | 2.4 (0-2.6) | 0 | 0.1 |
| G-CSF | 23.1 (0-50.4) | 0 (0-29) | 30.9 (19.3-50.4) | 0.1 |
| GM-CSF | 0.3 (0-0.9) | 0.5 (0-0.9) | 0.1 (0-0.4) | 0.4 |
| HGF | 468.5 (272.8-1900.4) | 1517  (453.8-1900.4) | 352.1 (272.8-483.1) | 0.1 |
| IFN-alpha* | - | - | - | - |
| IFN-gamma* | - | - | - | - |
| IL-1β | 2.6 (1.7-3.4) | 2.7 (2.6-3.4) | 2.2 (1.7-2.6) | 0.1 |
| IL-RA | 137.4 (118.3-1148.2) | 130.4  (118.3-1148.2) | 138.4 (136.4-587.5) | 0.5 |
| IL-2 | 1.8 (1.5-2.9) | 1.6 (1.5-2.9) | 2 (1.5-2.5) | 1.0 |
| IL-2R | 791.5 (396.1-2267.8) | 1069.2  (396.1-2267.8) | 513.7 (473-1124) | 0.8 |
| IL-4* | - | - | - | - |
| IL-5* | - | - | - | - |
| IL-6 | 11.4 (0.8-27.8) | 4.5 (0.8-18.3) | 25 (2.7-27.8) | 0.3 |
| IL-7* | - | - | - | - |
| IL-8 | 3.1 (0.9-12.2) | 2 (0.9-12.2) | 3.2 (3-5.2) | 0.5 |
| IL-10* | - | - | - | - |
| IL-12 | 170 (25.2-336.3) | 59.3  (25.2-336.3) | 246.7 (93-256.6) | 0.5 |
| IL-13 | 1.1 (0-1.9) | 1.7 (0.9-1.9) | 0.9 (0-1.2) | 0.1 |
| IL-15 | 2.3 (0-33.4) | 0 (0-33.4) | 3.9 (0.7-7.5) | 0.5 |
| IL-17A* | - | - | - | - |
| IP-10 | 10.3 (5.3-32) | 7.4 (5.3-32) | 13.2 (5.6-23.8) | 0.8 |
| MCP-1 | 74.6 (32-102.9) | 102.8 (32-102.9) | 74.6 (73-74.6) | 0.5 |
| MIG | 79.9 (49.5-170) | 65.8 (61.4-93.9) | 150.4 (49.5-170) | 0.5 |
| MIP-1α | - | - | - | - |
| MIP-1β | 15.3 (10.1-36.5) | 28.7 (16.3-36.5) | 10.8 (10-14.4) | **0.049** |
| RANTES | 4510.3 (114.6-8650) | 8650 (8650-8650) | 349.7 (114.6-370.6) | **0.04** |
| TNF-α | - | - | - | - |
| VEGF | 0 (0-2.9) | 0 (0-0.03) | 0 (0-2.9) | 0.8 |

***Supplementary Table 4*:** **Table showing differences in cytokine profiling at diagnosis between patients with chronic myelomonocytic leukemia with and without BM IDO1+ DC population (IDC). CMML plasma samples and time of collections overlapped with CyTOF.**

**Supplementary Figure 1: Supplementary figure 1 showing significant strong positive correlation between kynurenine concentration and T reg (%parent CD4 T) populations in plasma and PBMC samples collected at the same time-point and assessed through liquid-chromatography-mass spectrometry (LC-MS) and mass cytometry (CyTOF). Correlation was assessed via non-parametric Spearman test [rho (ρ) = 0.9, P=0.04*].**

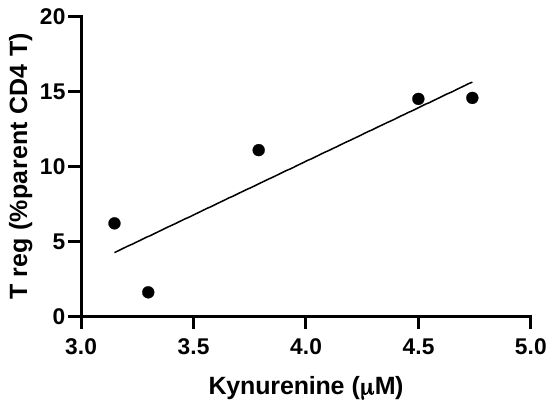

**Methods:**

**Immunohistochemistry and multiplex staining protocol:**

Immunohistochemical Staining:

Tissue sectioning and IHC staining was performed at the Pathology Research Core (Mayo Clinic, Rochester, MN) using the Leica Bond RX stainer (Leica). FFPE tissues were sectioned at 5 microns and IHC staining was performed on-line. Slides for IDO-1 stain were retrieved for 20 minutes using Epitope Retrieval 1 (Citrate; Leica) and incubated in Protein Block (Dako) for 5 minutes. The IDO-1 primary antibody (Rabbit Polyclonal, Sigma #HPA023072) was diluted to 1:300 in Background Reducing Diluent (Dako) and incubated for 15 minutes. The detection system used was Polymer Refine Detection System (Leica). This system includes the hydrogen peroxidase block, post primary and polymer reagent, DAB, and Hematoxylin. Immunostaining visualization was achieved by incubating slides 10 minutes in DAB and DAB buffer (1:19 mixture) from the Bond Polymer Refine Detection System. To this point, slides were rinsed between steps with 1X Bond Wash Buffer (Leica). Slides were counterstained for five minutes using Schmidt hematoxylin and molecular biology grade water (1:1 mixture), followed by several rinses in 1X Bond wash buffer and distilled water, this is not the hematoxylin provided with the Refine kit. Once the immunochemistry process was completed, slides were removed from the stainer and rinsed in tap water for five minutes. Slides were dehydrated in increasing concentrations of ethyl alcohol and cleared in 3 changes of xylene prior to permanent coverslipping in xylene-based medium. For CD11c and CD123 staining, clone 5D11 from Leica (Novocastra) and clone 7G3 from BD Pharmingen were used respectively.

Multiplex staining protocol:

Multiplex Immunofluorescence Staining and Visualization (CD11c, CD123, IDO-1 stains): Tissue sectioning and Immunofluorescence (IF) staining was performed at the Pathology Research Core (Mayo Clinic, Rochester, MN) using the Leica Bond RX stainer (Leica) and the Opal Polaris 7-Color Automation IHC Detection Kit (NEL871001KT; Akoya). This kit allows for the simultaneous detection of multiple proteins. Each Opal reactive fluorophore application is followed by heat-induced stripping steps before the application of the next antibody. The stripping steps remove the primary and secondary antibodies; which reduces any non-specific staining as well as tissue auto-fluorescence. Each staining cycle consist on incubating the slides in Akoya blocking buffer for five minutes, primary antibody incubation for 30 minutes, Opal Polymer HRP for 10 minutes, and Opal Fluorophore for 10 minutes. All primary antibodies were diluted in Akoya Antibody Diluent. Each Opal fluorophore used was first reconstituted in 75ul of DMSO, and then diluted in 1X Plus Automation Amplification Diluent for a final dilution of 1:150 prior to staining. FFPE tissues were sectioned at 5 microns. Slides were baked for 4 hours at 60°C then loaded in the instrument. Once in the stainer they were dewaxed and retrieved in Epitope Retrieval 2 (EDTA; Leica) for 20 minutes at 100°C. Slides were first stained with CD123 (1:200, mouse monoclonal; BD #554526), followed by Opal 570 fluorophore application. Primary and secondary antibodies were stripped using Epitope Retrieval 1 (Citrate; Leica) for 20 minutes at 95°C. Slides then were stained with IDO-1 (1:200, rabbit polyclonal; Sigma #HPA023072), followed by Opal 620 fluorophore application, and stripping steps. The last stain was CD11c (1:100, mouse monoclonal; Leica #NCL-L-CD11c-563), followed by Opal 690 fluorophore application. Counter stained was performed using Spectral DAPI (Akoya) for five minutes. One additional slide was process following the same steps but without any of the Opal fluorophores or the Spectral DAPI. This slide is use to determine the autofluorescence signal of the tissue so it can be spectrally remove from the final images during the unmixing process perform within the Imaging system. Once the IF process was completed, slides were removed from the stainer and rinsed in tap water for three minutes. Slides were perminatly coverslipped using glycerol-based liquid mountant (Prolong Diamond Antifade Mountant; Invitrogen). Tissues were scanned using the MOTif technology within the Vectra Polaris Automated Quantitative Pathology Imaging System (Akoya). Once scanned, areas were selected for unmixing using the Phenochart software, and image unmixing was performed using InForm software. Due to lack of previous papers and the novelty of the staining technology and imaging this methods sections was produced based on information from the supplier.

**Quantification of Tryptophan and Kynurenine metabolites**

**(Liquid-Chromatography-Mass Spectrometry):**

Quantitative analysis of tryptophan and its resultant kynurenine metabolites were performed by the Mayo Clinic Metabolomics Core as previously described by Lanza IR et al. *PLoS One* 2010;5(5):e10538, and summarized as follows. Plasma samples and calibration standards were prepared with MassTrak Amino Acid Analysis Solution (AAA) kit from Waters according to instructions with slight modifications for detection on a mass spectrometer. A 10-point standard concentration curve was made from the calibration standard solution to calculate tryptophan and its resultant kynurenine metabolites concentrations in plasma samples. A solution containing the metabolite isotopes of interest were purchased from Cambridge Isotope Laboratories, Isotec and MassTrace was used as the internal standard solution. Frozen plasma samples were thawed, spiked with internal standard then deproteinized with cold MeOH followed by centrifugation at 10,000 g for 5 minutes prior to derivatization with the derivatizing reagent 6-aminoquinolyl-N-hydroxysuccinimidyl carbamate according to MassTrak instructions. High resolution separation was done using an Acquity UPLC system, injecting 1 µl of derivatized solution, with a UPLC BEH C18 1.7 micron 2.1×150 mm column from Waters. Column flow was set to 400 µl/min with a gradient from 99.9%A to 98%B where buffer A is 1% acetonitrile in 0.1% formic acid and buffer B is 100% acetonitrile A column temp of 43 degrees Celsius and a sample tray temp of 6% Celsius was set and mass detection was completed on a TSQ Quantum Ultra from Thermo Finnigan running in positive ESI mode. Finally, the settings of a scan width of 0.002, scan time of 0.04 seconds per transition mass, collision energy of 25, collision gas pressure of 1.5 mTorr, tube lens value set to 90 and monitoring a signature ion of the derivatized amines at m/z 171.04 by selected reaction monitoring were employed. By using these aforementioned methods, tryptophan and its resultant kynurenine metabolites were able to be measured.

**Mass Cytometry (CyTOF) Protocol:**

*Antibody and sample preparation:* For primary conjugations, purified antibodies were obtained in carrier protein-free PBS and labeled using the X8 polymer MaxPAR antibody conjugation kit (Fluidigm) according to the manufacturer’s protocol. The antibody panel was designed using the web-based Fluidigm panel designer to select channels with optimal signal and minimal background from oxidation, isotopic impurity or abundance sensitivity. All antibodies were titrated to optimal staining concentrations using primary human bone marrows of patients with MM. Antibody master mixes were prepared fresh for each experiment. All BM and PB samples were processed identically. Mononuclear cells were obtained after ACK lysis of BM and PB samples and were viably frozen in RPMI 1640, 20%FBS, 10% DMSO. Cryopreserved cells were resuscitated for mass cytometry analyses by rapid thawing and were rested in RPMI 1640 (20% FBS) for 60 minutes prior to staining. Staining was performed using Fluidigm’s protocol. Briefly, 1-3 million cells were stained for viability with 5mM cisplatin for 5 mins at room temperature and quenched with cell staining medium (CSM, Fluidigm). Cells were then incubated for 10 mins at room temperature with human FcR blocking reagent (Biolegend) and then stained with the surface antibody cocktail for 60mins at 4^o^C with gentle agitation. Finally, cells were washed twice with CSM, fixed with 1.6% PFA, washed with CSM and resuspended in 1:1000 solution of Iridium intercalator diluted in MaxPar Fix and Perm buffer (Fluidigm) for 20mins at room temperature. Prior to acquisition, cells were washed twice in CSM and twice in deionized water and were then diluted to a concentration 0.5million cells/ml in water containing 10% of EQ 4 Element Beads (Fluidigm). Cells were filtered through a 35μm membrane prior to mass cytometry acquisition. Samples were then acquired on a Helios mass cytometer.

*CyTOF panel details:*

| ***No.*** | ***Product Code*** | ***Name*** | ***Species*** | ***Clone*** | ***Metal*** | ***Target*** | ***Cell*** |
| --- | --- | --- | --- | --- | --- | --- | --- |
| 1 | 3089003B | Anti-Human CD45 (HI30)-Y89—100 Tests | Human | HI30 | 89Y | CD45 | Lymphocytes |
| 2 | 3141003A | Anti-Human CD196/CCR6 (G034E3)-141Pr—50 Tests | Human | G034E3 | 141Pr | CD196/CCR6 | T-cell - Th17 |
| 3 | 3142001B | Anti-Human CD19 (HIB19)-142Nd—100 Tests | Human | HIB19 | 142Nd | CD19 | B-cell |
| 4 | 3143012B | Anti-Human CD127/IL-7Ra (A019D5)-143Nd—100 Tests | Human | A019D5 | 143Nd | CD127/IL-7Ra | T-cell - Memory |
| 5 | 3144014B | Anti-Human CD38 (HIT2)-144Nd—100 Tests | Human | HIT2 | 144Nd | CD38 | Many - Activation |
| 6 | 3146005B | Anti-Human IgD (IA6-2)-146Nd—100 Tests | Human | IA6-2 | 146Nd | IgD | B-cell |
| 7 | 3147008B | Anti-Human CD11c (Bu15)-147Sm—100 Tests | Human | Bu15 | 147Sm | CD11c | Dendritic Cells |
| 8 | 3148004B | Anti-Human CD16 (3G8)-148Nd—100 Tests | Human | 3G8 | 148Nd | CD16 | NK cells |
| 9 | 3149029C | Anti-Human CD194/CCR4 (205410)-149Sm—50 Tests | Human | L291h4 | 149Sm | CD194/CCR4 | T-cell - Th2 |
| 10 | 3151001B | Anti-Human CD123/IL-3R (6H6)-151Eu—100 Tests | Human | 6H6 | 151Eu | CD123/IL-3R | Myeloid cells |
| 11 | 3152008B | Anti-Human TCRgd (11F2)-152Sm—100 Tests | Human | 11F2 | 152Sm | TCRgd | g/d T-cell |
| 12 | 3153020B | Anti-Human CD185/CXCR5 (RF8B2)-153Eu—100 Tests | Human | RF8B2 | 153Eu | CD185/CXCR5 | B-cell |
| 13 | 3154003B | Anti-Human CD3 (UCHT1)-154Sm—100 Tests | Human | UCHT1 | 154Sm | CD3 | T-cell |
| 14 | 3155011B | Anti-Human CD45RA (HI100)-155Gd—100 Tests | Human | HI100 | 155Gd | CD45RA | T-cell - Naïve |
| 15 | 3158010B | Anti-Human CD27 (L128)-158Gd—100 Tests | Human | L128 | 158Gd | CD27 | B-cell - Memory; T-cell |
| 16 | 3160003B | Anti-Human CD28 (CD28.2)-160Gd—100 Tests | Human | CD28.2 | 160Gd | CD28 | T-cell |
| 17 | 3162023B | Anti-Human CD66b (80H3)-162Dy—100 Tests | Human | 80H3 | 162Dy | CD66b | Granulocytes |
| 18 | 3163004B | Anti-Human CD183/CXCR3 (G025H7)-163Dy—100 Tests | Human | G025H7 | 163Dy | CD183/CXCR3 | T-cell - Th1 |
| 19 | 3164009B | Anti-Human CD161 (HP-3G10)-164Dy—100 Tests | Human | HP-3G10 | 164Dy | CD161 | NK cells |
| 20 | 3165011B | Anti-Human CD45RO (UCHL1)-165Ho—100 Tests | Human | UCHL1 | 165Ho | CD45RO | T-cell - Memory |
| 21 | 3166007B | Anti-Human CD24 (ML5)-166Er—100 Tests | Human | ML5 | 166Er | CD24 | B-cell |
| 22 | 3167009A | Anti-Human CD197/CCR7 (G043H7)-167Er—50 Tests | Human | G043H7 | 167Er | CD197/CCR7 | T-cell - Naïve/Memory |
| 23 | 3168002B | Anti-Human CD8 (SK1)-168Er—100 Tests | Human | SK1 | 168Er | CD8a | T-cell |
| 24 | 3169003B | Anti-Human CD25 (2A3)-169Tm—100 Tests | Human | 2A3 | 169Tm | CD25/IL-2R | T-cell - T-reg |
| 25 | 3171012B | Anti-Human CD20 (2H7)-171Yb—100 Tests | Human | 2H7 | 171Yb | CD20 | B-cell |
| 26 | 3172009B | Anti-Human CD57 (HCD57)-172Yb—100 Tests | Human | HCD57 | 172Yb | CD57 | NK Cells |
| 27 | 3173005B | Anti-Human HLA-DR (L243)-173Yb—100 Tests | Human | L243 | 173Yb | HLA-DR | DC, Mono, Macs |
| 28 | 3174004B | Anti-Human CD4 (SK3)-174Yb—100 Tests | Human | SK3 | 174Yb | CD4 | T-cell |
| 29 | 3175015B | Anti-Human CD14 (M5E2)-175Lu—100 Tests | Human | M5E2 | 175Lu | CD14 | Monocytes |
| 30 | 3176008B | Anti-Human CD56 (NCAM16.2)-176Yb—100 Tests | Human | NCAM16.2 | 176Yb | CD56/NCAM | T-cells - Activation |

**Cytokine analysis**

Plasma samples from CMML cases and normal controls were tested for the presence of 30 cytokines including: G-CSF, GM-CSF, Eotaxin, IP-10, EGF, FGF-basic, IFN-gamma, IFN-alpha, IL-1beta, IL-1RA, MCP-1, MIG, MIP-1alpha, MIP-1beta, HGF, VEGF, IL-10, IL-2, IL-2R, IL-4, IL-5, IL-6, IL-7, IL-8, IL-12, IL-13, IL-15, IL-17, RANTES, and TNF-alpha, by the Cytokine Human Magnetic 30-Plex Panel Kit (ThermoFisher Scientific, San Francisco, CA). The bead mix was prepared according to the manufacturer’s protocol and samples were analyzed in a multiplex array using ProcartaPlex magnetic beads via a Luminex^®^ 200™ instrument (Austin, TX). All plasma samples were run in duplicate. The data were analyzed using the instrument specific software, xPONENT®.

**Whole transcriptome shotgun sequencing (RNA-seq) protocol and analysis**

Library preparation for the RNA samples was done using Illumina TruSeq Stranded Total RNA and sequencing was done using Illumina HiSeq 2500, 100 cycles x 2 paired-end reads. RNA-seq sequencing reads were processed through the MAPRSeq v2.0. bioinformatics workflow as described in (Kalari et al., 2014). Briefly, reads were mapped using TopHat version 2.06 against the reference GRCh37 without alternative haplotypes using the transcript models from UCSD (March 2012) available from Illumina iGenomes Project. Gene counts were calculated with featureCounts (Liao et al., 2014). RNA samples were prepared and RNA quality was determined using an Agilent Bioanalyzer RNA Nanochip or Caliper RNA assay and arrayed into a 96-well plate. The paired-end sequencing libraries were prepared following BC Cancer Genome Sciences Centre’s strand-specific, plate-based library construction protocol on a Microlab NIMBUS robot (Hamilton Robotics, USA). Libraries were sequenced on an Illumina HiSeq2500 platform to a read depth of approximately 50 million reads per sample. Patient samples were grouped based on the previously identified histological classifications and gene set enrichment analysis (Subramanian, Tamayo, et al. (2005), PNAS 102, 15545-15550, http://www.broad.mit.edu/gsea/) was carried out in untreated patients from two groups: CMML patients with BM IDO1+ dendritic cell populations (IDC, n=4) and CMML patients without BM IDC (n=4).

**Statistical analysis**

Distribution of continuous variables was statistically compared using non-parametric (Mann-Whitney or Kruskal-Wallis) tests, while nominal variables were compared using the Chi-Square test. Through Kaplan-Meier method, overall survival (OS) was computed from date of diagnosis to date of death or last follow-up, while acute myeloid leukemia (AML)-free survival (LFS) was computed from date of diagnosis to the date of AML transformation (AML transformation replaced death as the censored event for LFS). Patients treated with allogeneic hematopoietic stem cell transplant were censored at the time of transplant for both OS and LFS. Statistics were performed via Graphpad Prism (version 8.1.2) and JMP (version 14.1.0) softwares and a *P* value of ≤ 0.05 was considered statistically significant.
